# The architecture of SARS-CoV-2 transcriptome

**DOI:** 10.1101/2020.03.12.988865

**Authors:** Dongwan Kim, Joo-Yeon Lee, Jeong-Sun Yang, Jun Won Kim, V. Narry Kim, Hyeshik Chang

## Abstract

SARS-CoV-2 is a betacoronavirus that is responsible for the COVID-19 pandemic. The genome of SARS-CoV-2 was reported recently, but its transcriptomic architecture is unknown. Utilizing two complementary sequencing techniques, we here present a high-resolution map of the SARS-CoV-2 transcriptome and epitranscriptome. DNA nanoball sequencing shows that the transcriptome is highly complex owing to numerous recombination events, both canonical and noncanonical. In addition to the genomic RNA and subgenomic RNAs common in all coronaviruses, SARS-CoV-2 produces a large number of transcripts encoding unknown ORFs with fusion, deletion, and/or frameshift. Using nanopore direct RNA sequencing, we further find at least 41 RNA modification sites on viral transcripts, with the most frequent motif being AAGAA. Modified RNAs have shorter poly(A) tails than unmodified RNAs, suggesting a link between the internal modification and the 3′ tail. Functional investigation of the unknown ORFs and RNA modifications discovered in this study will open new directions to our understanding of the life cycle and pathogenicity of SARS-CoV-2.

**Highlights:**

- We provide a high-resolution map of SARS-CoV-2 transcriptome and epitranscriptome using nanopore direct RNA sequencing and DNA nanoball sequencing.
- The transcriptome is highly complex owing to numerous recombination events, both canonical and noncanonical.
- In addition to the genomic and subgenomic RNAs common in all coronaviruses, SARS-CoV-2 produces transcripts encoding unknown ORFs.
- We discover at least 41 potential RNA modification sites with an AAGAA motif.

## Introduction

Coronavirus disease 19 (COVID-19) is caused by a novel coronavirus designated as severe acute respiratory syndrome coronavirus 2 (SARS-CoV-2) (Zhou et al., 2020; Zhu et al., 2020). Like other coronaviruses (order *Nidovirales*, family *Coronaviridae*, subfamily *Coronavirinae*), SARS-CoV-2 is an enveloped virus with a positive-sense, single-stranded RNA genome of ~30 kb. SARS-CoV-2 belongs to the genus *betacoronavirus*, together with SARS-CoV and Middle East respiratory syndrome coronavirus (MERS-CoV) (with 80% and 50% homology, respectively) (Kim et al., 2020; Zhou et al., 2020). Coronaviruses (CoVs) were thought to primarily cause enzootic infections in birds and mammals. But, the recurring outbreaks of SARS, MERS, and now COVID-19 have clearly demonstrated the remarkable ability of CoVs to cross species barriers and transmit between humans (Menachery et al., 2017).

CoVs carry the largest genomes (26-32 kb) among all RNA virus families (Figure 1). Each viral transcript has a 5′-cap structure and a 3′ poly(A) tail (Lai and Stohlman, 1981; Yogo et al., 1977). Upon cell entry, the genomic RNA is translated to produce nonstructural proteins (nsps) from two open reading frames (ORFs), ORF1a, and ORF1b. The ORF1a produces polypeptide 1a (pp1a, 440-500 kDa) that is cleaved into 11 nsps. The –1 ribosome frameshift occurs immediately upstream of the ORF1a stop codon, which allows continued translation of ORF1b, yielding a large polypeptide (pp1ab, 740-810 kDa) which is cleaved into 16 nsps. The proteolytic cleavage is mediated by viral proteases nsp3 and nsp5 that harbor a papain-like protease domain and a 3C-like protease domain, respectively.

**Figure 1.**
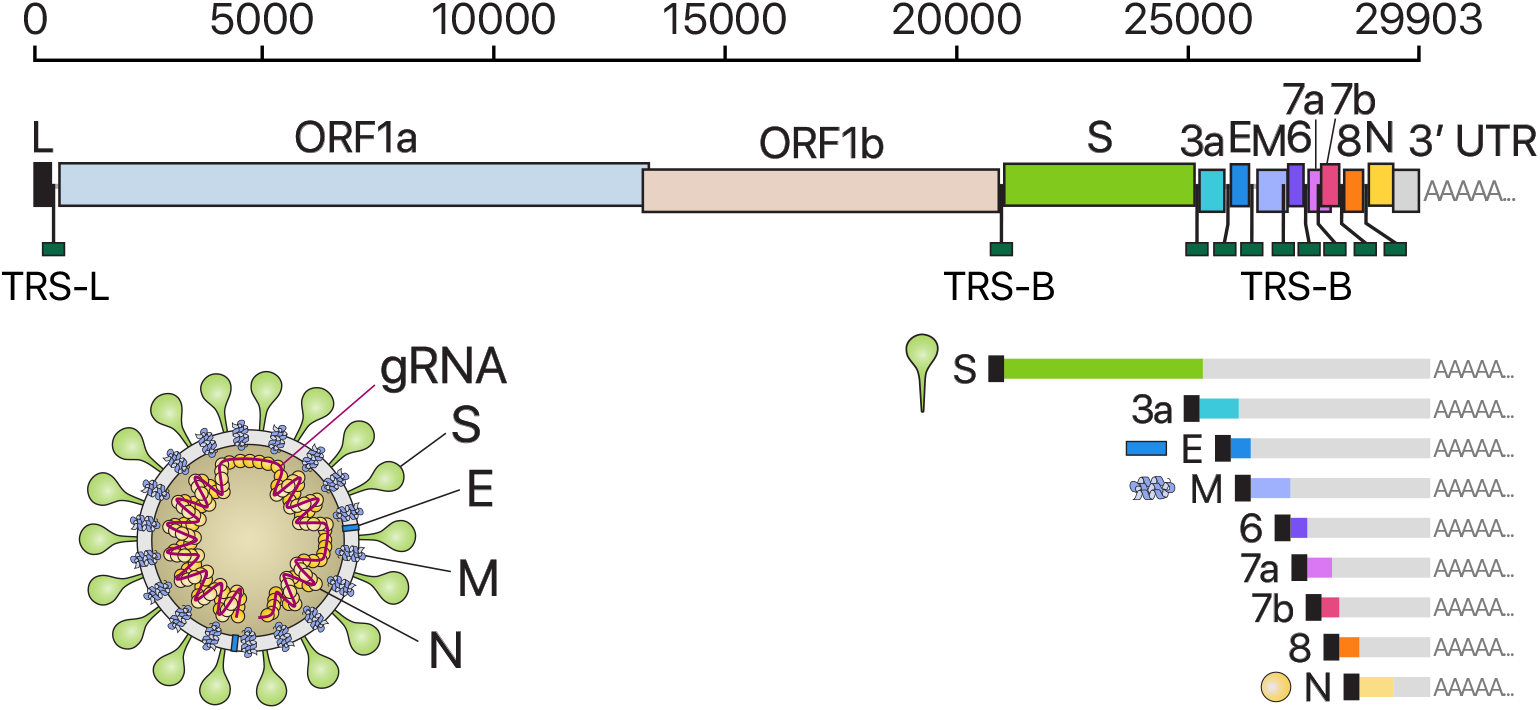
Schematic presentation of the SARS-CoV-2 genome organization, the canonical subgenomic mRNAs, and the virion structure. From the full-length genomic RNA (29,903 nt) which also serves as an mRNA, ORF1a and ORF1b are translated. In addition to the genomic RNA, nine major subgenomic RNAs are produced. The sizes of the boxes representing small accessory proteins are bigger than the actual size of the ORF for better visualization. The black box indicates leader sequence. Note that ORF10 is not included here because our data show no evidence for ORF10 expression.

The viral genome is also used as the template for replication and transcription, which is mediated by nsp12 harboring RNA-dependent RNA polymerase (RdRP) activity (Snijder et al., 2016; Sola et al., 2015). Negative-strand RNA intermediates are generated to serve as the templates for the synthesis of positive-sense genomic RNA (gRNA) and subgenomic RNAs (sgRNAs). The gRNA is packaged by the structural proteins to assemble progeny virions. Shorter sgRNAs encode conserved structural proteins (spike protein (S), envelope protein (E), membrane protein (M), and nucleocapsid protein (N)), and several accessory proteins. SARS-CoV-2 is known to have six accessory proteins (3a, 6, 7a, 7b, 8, and 10) according to the current annotation (NCBI Reference Sequence: NC_045512.2). But the ORFs have not yet been experimentally verified for expression. Therefore, it is currently unclear which accessory genes are actually expressed from this compact genome.

Each coronaviral RNA contains the common 5′ “leader” sequence of 70-100 nt fused to the “body” sequence from the 3′ end of the genome (Lai and Stohlman, 1981; Sola et al., 2015) (Figure 1). According to the prevailing model, leader-to-body fusion occurs during negative-strand synthesis at short motifs called transcription-regulating sequences (TRSs) that are located immediately adjacent to ORFs (Figure 1). TRSs contain a conserved 6-7 nt core sequence (CS) surrounded by variable sequences. During negative-strand synthesis, RdRP pauses when it crosses a TRS in the body (TRS-B), and switches the template to the TRS in the leader (TRS-L), which results in the leader-body fusion. From the fused negative-strand intermediates, positive-strand mRNAs are transcribed. The replication and transcription mechanism has been studied in other coronaviruses. However, it is unclear whether the general mechanism also applies to SARS-CoV-2 and if there are any unknown components in the SARS-CoV-2 transcriptome. For the development of diagnostic and therapeutic tools and the understanding of this new virus, it is critical to define the organization of the SARS-CoV-2 genome.

Deep sequencing technologies offer a powerful means to investigate viral transcriptome. The “sequencing-by-synthesis (SBS)” methods such as the Illumina and MGI platforms confer high accuracy and coverage. But they are limited by short read length (200-400 nt), so the fragmented sequences should be re-assembled computationally, during which the haplotype information is lost. More recently introduced is the nanopore-based direct RNA sequencing (DRS) approach. While nanopore DRS is limited in sequencing accuracy, it enables long-read sequencing, which would be particularly useful for the analysis of long nested CoV transcripts. Moreover, because DRS does not require reverse transcription to generate cDNA, the RNA modification information can be detected directly during sequencing. Numerous RNA modifications have been found to control eukaryotic RNAs and viral RNAs (Williams et al., 2019). Terminal RNA modifications such as RNA tailing also plays a critical role in cellular and viral RNA regulation (Warkocki et al., 2018).

In this study, we combined two complementary sequencing approaches, DRS and SBS. We unambiguously mapped the sgRNAs, ORFs, and TRSs of SARS-CoV-2. Additionally, we found numerous unconventional recombination events that are distinct from canonical TRS-mediated joining. We further discovered RNA modification sites and measured the poly(A) tail length of gRNAs and sgRNAs.

## Results and Discussion

To delineate the SARS-CoV-2 transcriptome, we first performed DRS runs on a MinION nanopore sequencer using total RNA extracted from Vero cells infected with SARS-CoV-2 (BetaCoV/Korea/KCDC03/2020). The virus was isolated from a patient who was diagnosed with COVID-19 on January 26, 2020, after traveling from Wuhan, China (Kim et al., 2020). We obtained 879,679 reads from infected cells (corresponding to a throughput of 1.9 Gb) (Figure 2A). The majority (65.4%) of the reads mapped to SARS-CoV-2, indicating that viral transcripts dominate the transcriptome while the host gene expression is strongly suppressed. Although nanopore DRS has the 3′ bias due to directional sequencing from the 3′-ends of RNAs, approximately half of the viral reads still contained the 5′ leader.

**Figure 2.**
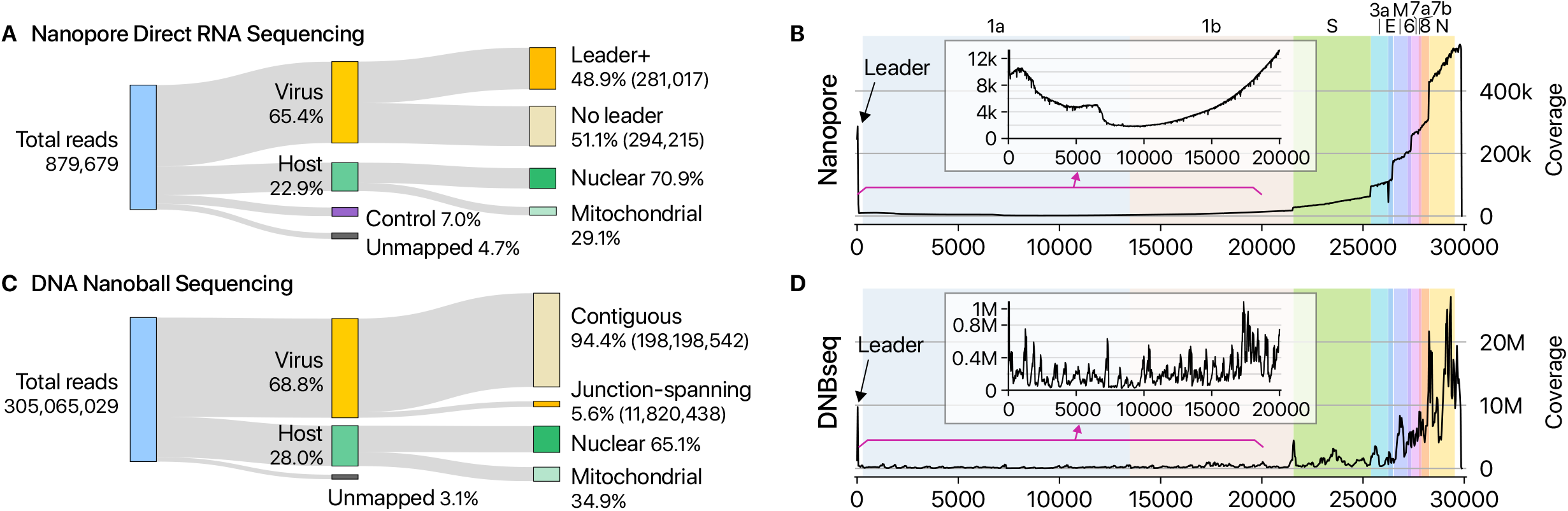
Statistics of sequencing data. **A,** Read counts from nanopore direct RNA sequencing of total RNA from Vero cells infected with SARS-CoV-2. “Leader+” indicates the viral reads that contain the *5′* end leader sequence. “No leader” denotes the viral reads lacking the leader sequence. “Nuclear” reads match to mRNAs from the nuclear chromosome while “mitochondrial” reads are derived from the mitochondrial genome. “Control” indicates quality control RNA for nanopore sequencing. **B,** Genome coverage of the nanopore direct RNA sequencing data shown in panel A. The stepwise reduction in coverage corresponds to the borders expected for the canonical sgRNAs. The smaller inner plot magnifies the 5′ part of the genome. **C,** Read counts from DNA nanoball sequencing using MGISEQ. Total RNA from Vero cells infected with SARS-CoV-2 was used for sequencing. **D,** Genome coverage of the DNA nanoball sequencing (DNBseq) data shown in panel C.

The SARS-CoV-2 genome was fully covered, missing only 12 nt from the 5′ end (Figure 2B). The longest tags (111 reads) correspond to the full-length gRNA (Figure 2B). The coverage of the 3′ side of the viral genome is substantially higher than that of the 5′ side, which reflects the nested sgRNAs. This is also partly due to the 3′ bias of the directional DRS technique. The presence of the leader sequence (72 nt) in viral RNAs results in a prominent coverage peak at the 5′ end, as expected. We could also clearly detect vertical drops in the coverage, whose positions correspond to the leader-body junction in sgRNAs. All known sgRNAs are supported by DRS reads, with the exception of ORF10 (see below).

In addition, we observed unexpected reads reflecting noncanonical recombination events. Such fusion transcripts resulted in the increased coverage towards the 5′ end (Figure 2B, inner box). Early studies on coronavirus mouse hepatitis virus reported that recombination frequently occurs (Furuya and Lai, 1993; Liao and Lai, 1992; Luytjes et al., 1996). Some viral RNAs contain the 5′ and 3^’^ proximal sequences resulting from “illegitimate” recombination events.

To further validate sgRNAs and their junction sites, we performed DNA nanoball sequencing based on the sequencing-by-synthesis principle (DNBseq) and obtained 305,065,029 reads (Figure 2C). The results are overall consistent with the DRS data. The leader-body junctions are frequently sequenced, giving rise to a sharp peak at the 5′ end in the coverage plot (Figure 2D). The 3′ end exhibits a high coverage as expected for the nested transcripts.

The depth of DNB sequencing allowed us to confirm and examine the junctions on an unprecedented scale for a CoV genome. We mapped the 5′ and 3′ sites at the fused junctions and estimated the recombination frequency by counting the reads spanning the junctions (Figure 3A). The leader represents the most prominent 5′ site, as expected (Figure 3A, red asterisk on the x-axis). The known TRSs are detected as the top 3′ sites (Figure 3A, red dots on the y-axis). These results confirm that SARS-CoV-2 uses the canonical TRS-mediated mechanism for discontinuous transcription to produce major sgRNAs (Figure 3B). Quantitative comparison of the junction-spanning reads shows that the N RNA is the most abundantly expressed transcript, followed by S, 7a, 3a, 8, M, E, 6, and 7b (Figure 3C). It is important to note that ORF10 is represented by only one read in DNB data (0.000009 % of viral junction-spanning reads) and that it was not supported at all by DRS data. ORF10 does not show significant homology to known proteins. Thus, ORF10 is unlikely to be expressed, and the annotation of ORF10 should be reconsidered. Taken together, SARS-CoV-2 expresses nine canonical sgRNAs (S, 3a, E, M, 6, 7a, 7b, 8, and N) together with the gRNA (Figures 1 and 3C).

**Figure 3.**
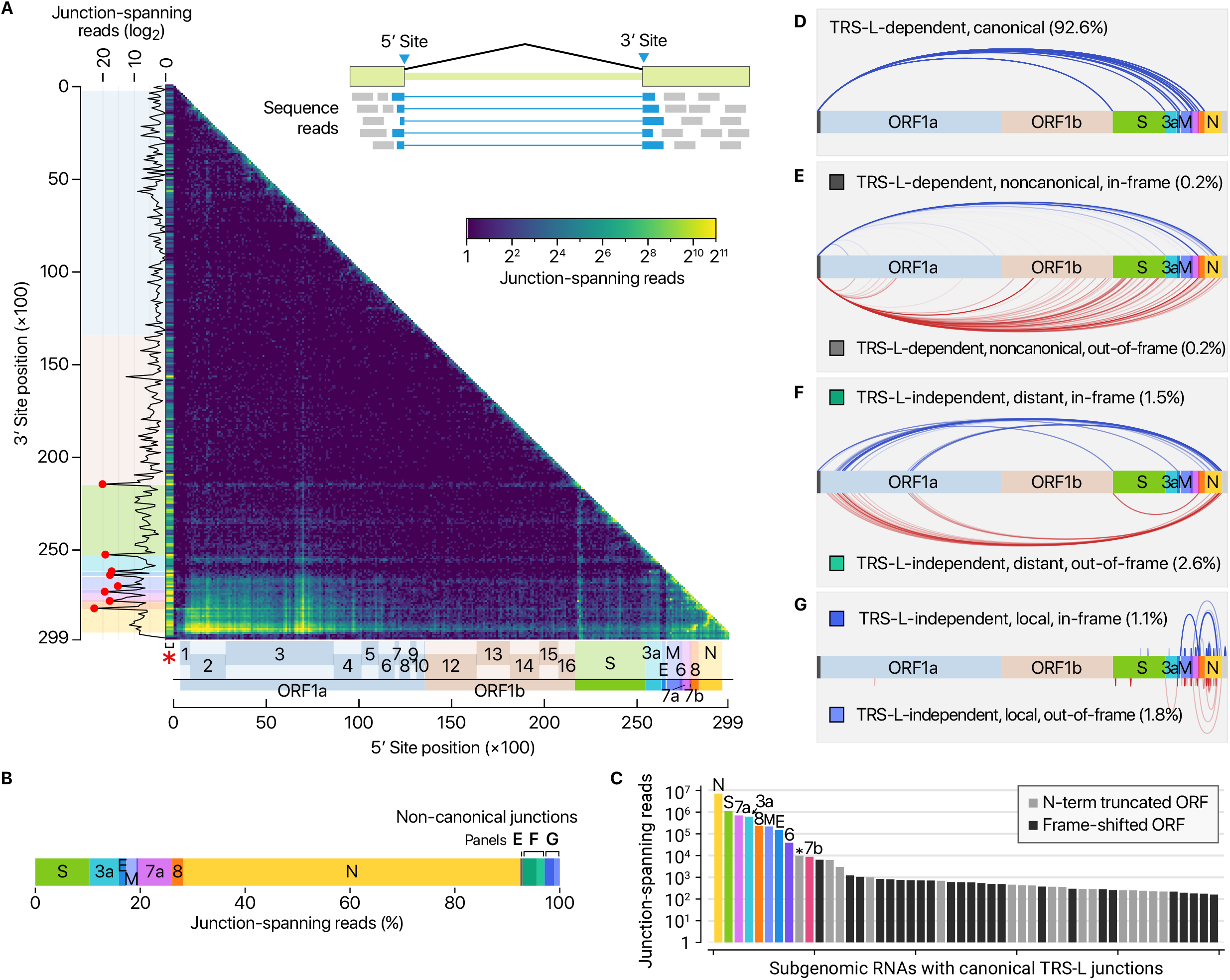
Viral subgenomic RNAs and their recombination sites. **A,** Positions of recombination sites determined by using junction-spanning reads from DNBseq data. The x-axis and y-axis show the positions of the 5′ sites and their joined 3′ sites, respectively. The frequency of the recombination was estimated by the read counts of the junction-spanning reads. The red asterisk on the x-axis is to indicate that the leader sequence. Please note that the left-most bins containing the leader TRS was expanded horizontally on this heatmap to improve visualization. The red dots on the sub-plot alongside the y-axis denote local peaks which coincide with the 5′ end of the body of each sgRNA. **B,** Transcript abundance was estimated by counting the DNBseq reads that span the junction of the corresponding RNA. **C,** Top 50 sgRNAs. The asterisk indicates an ORF beginning at 27,825 which may encode the 7b protein with an N-terminal truncation of 23 amino acid. The grey bars denote minor transcripts that encode proteins with an N-terminal truncation compared with the corresponding overlapping transcript. The black bars indicate minor transcripts that encode proteins in a different reading frame from the overlapping major mRNA. **D,** Canonical recombination. **E,** TRS-L-dependent noncanonical recombination between the leader TRS and a noncanonical 3′ site in the body. **F,** TRS-L-independent long-distance (>5,000 nt) recombination. **G,** TRS-L-independent local recombination yielding a deletion between proximal sites (20-5,000 nt distance).

In addition to the canonical sgRNAs with expected structure and length (Figure 3D), our results show many minor recombination sites (Figures 3E-3G, Table S2). There are three main types of such recombinant events. The RNAs in the first cluster have the leader combined with the body at unexpected 3′ sites in the middle of ORFs or UTRs (Figure 3E, TRS-L-dependent noncanonical recombination; Table S3). The second cluster shows a long-distance fusion between sequences that do not have similarity to the leader (Figure 3F, TRS-L-independent distant recombination). The last group undergoes local recombination, which leads to smaller deletions, mainly in structural and accessory genes, including S (Figure 3G, TRS-L-independent local recombination).

Of note, the junctions in these noncanonical transcripts do not contain a known TRS-B, indicating that at least some of these transcripts may be generated through a different mechanism(s). It was previously shown in other coronaviruses that transcripts with partial sequences are produced (Furuya and Lai, 1993; Liao and Lai, 1992; Luytjes et al., 1996). These RNAs are considered as parasites that compete for viral proteins, hence referred to as “defective interfering RNAs” (DI-RNAs) (Pathak and Nagy, 2009). Similar sgRNAs have also been described in a recent sequencing analysis on alphacoronavirus HCoV-229E (Viehweger et al., 2019), suggesting this mechanism may be conserved among coronaviruses. While this may be due to erroneous replicase activity, it remains an open question if the noncanonical recombinations have an active role in the viral life cycle and evolution. Although individual RNA species are not abundant, the combined read numbers are often comparable to the levels of accessory transcripts. Most of the RNAs have coding potential to yield proteins. Many transcripts (that belong to the “TRS-L-independent distant” group) encode the upstream part of ORF1a, including nsp1, nsp2, and truncated nsp3, which may change the stoichiometry between nsps (Figure 3F, Table S4). Another notable example is the 7b protein with an N-terminal truncation that may be produced at a level similar to the annotated full-length 7b (Figure 3C, asterisk). Frame-shifted ORFs may also generate short peptides that are different from known viral proteins (Figure 3B). It will be interesting in the future to examine if these unknown ORFs are actually translated and yield functional products.

As nanopore DRS is based on single-molecule detection of RNA, it offers a unique opportunity to examine multiple epitranscriptomic features of individual RNA molecules. We recently developed software to measure the length of poly(A) tail from DRS data (Choi and Chang, unpublished). Using this software, we confirm that, like other CoVs, SARS-CoV-2 RNAs carry poly(A) tails (Figures 4A-B). The tail is likely to be critical for both translation and replication. The tail of viral RNAs is 47 nt in median length. The full-length viral RNA has a relatively longer tail than sgRNAs. Notably, sgRNAs have two tail populations: a minor peak at ~30 nt and a major peak at ~45 nt (Figure 4B, arrowheads). Wu and colleagues previously observed that the poly(A) tail length of bovine CoV mRNAs change during infection: from ~45 nt immediately after virus entry to ~65 nt at 6-9 h.p.i. and ~30 nt at 120-144 h.p.i. (Wu et al., 2013). Thus, the short tails of ~30 nt observed in this study may represent aged RNAs that are prone to decay. Viral RNAs exhibit a homogenous length distribution, unlike host nuclear genome-encoded mRNAs (Figure 4C). The distribution is similar to that of mitochondrial chromosome-encoded RNAs whose tail is generated by MTPAP (Tomecki et al., 2004). It was recently shown that HCoV-229E nsp8 has an adenylyltransferase activity, which may extend poly(A) tail of viral RNA (Tvarogova et al., 2019). Because poly(A) tail should be constantly attacked by host deadenylases, the regulation of viral RNA tailing is likely to be important for viral end replication.

**Figure 4.**
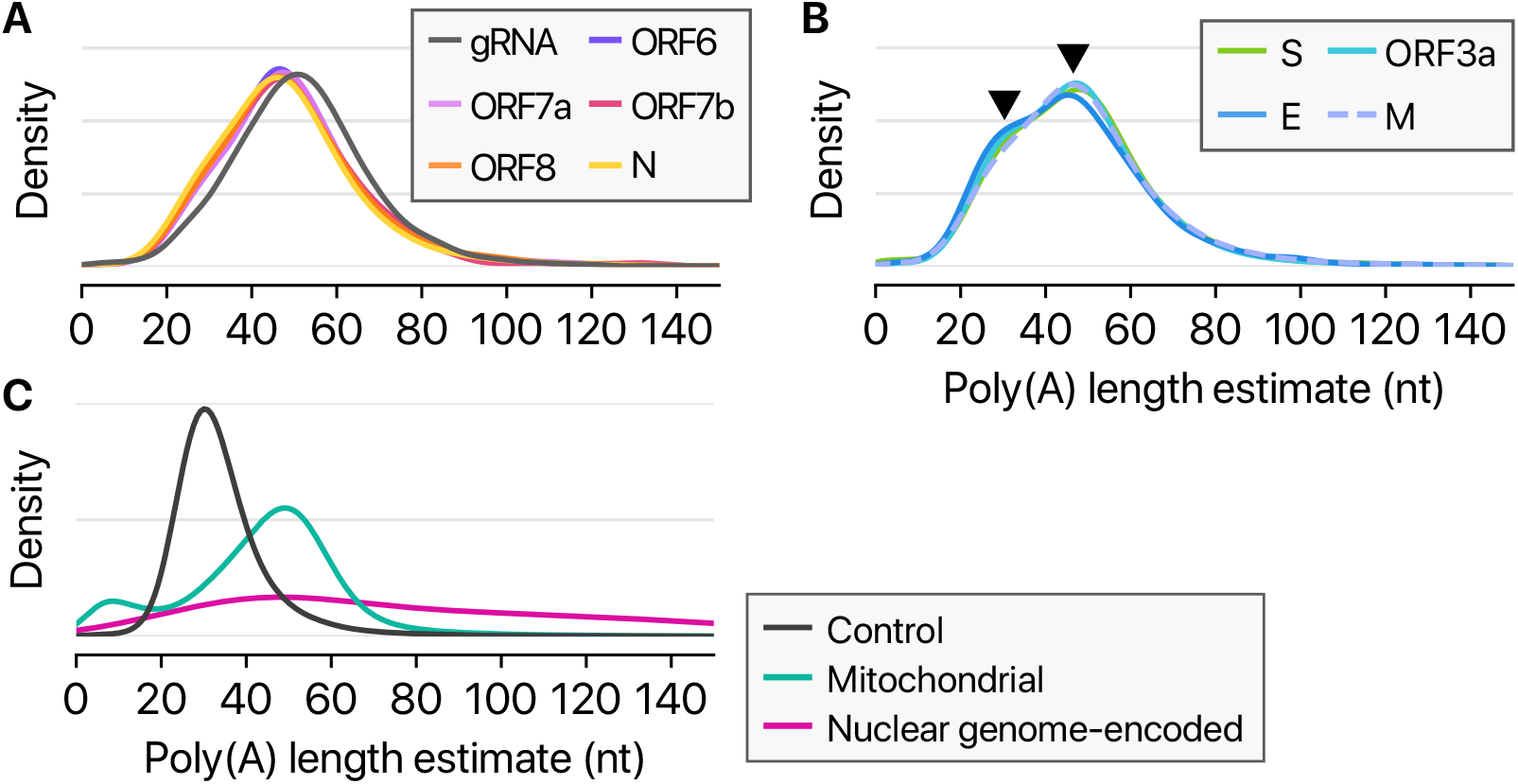
Length of poly(A) tail. **A-B,** Poly(A) tail length distribution of viral transcripts. Arrowheads indicate peaks at ~30 and ~45 nt. **C,** Poly(A) tail length distribution of quality control RNA which has a 30-nt poly(A) tail, host mRNAs from the nuclear chromosome, or host RNAs from the mitochondrial chromosome.

Next, we examined the epitranscriptomic landscape of SARS-CoV-2 by using the DRS data. Viral RNA modification was first described more than 40 years ago (Gokhale and Horner, 2017). N6-methyladenosine (m6A) is the most widely used modification (Courtney et al., 2017; Gokhale et al., 2016; Krug et al., 1976; Lichinchi et al., 2016; Narayan et al., 1987), but other modifications have also been reported on viral RNAs, including 5-methylcytosine methylation (5mC), 2′-O-methylation (Nm), deamination, and terminal uridylation. In a recent analysis of HCoV-229E using DRS, modification calling suggested frequent 5mC signal across viral RNAs (Viehweger et al., 2019). But since no direct control group was included in the analysis, the proposed modification needed validation. To unambiguously investigate the modifications, we generated negative control RNAs by in vitro transcription of the viral sequences and performed a DRS run on these unmodified controls (Figure S1A). The partially overlapping control RNAs are ~2.3 kb or ~4.4 kb each and cover the entire length of the genome (Figure S1B). Detection using pre-trained models reported numerous signal level changes corresponding to 5mC modification, even with the unmodified controls (Figure S1C). We obtained highly comparable results from the viral RNAs from infected cells (Figure S1D), demonstrating that the 5mC sites detected without a control may be false positives.

We, however, noticed intriguing differences in the ionic current (called “squiggles”) between negative control and viral transcripts (Figure 5A). At least 41 sites displayed substantial differences (over 20% frequency), indicating potential RNA modifications (Table S5). Notably, some of the sites showed different frequencies depending on the sgRNA species. Figures 5A-5C show an example that is modified more heavily on the S RNA than the N RNA, while Figures S2A-S2C present a site that is modified more frequently on the ORF8 RNA compared with the S RNA. Moreover, the dwell time of the modified base is longer than that of the unmodified base (Figure 5D, right), suggesting that the modification interferes with the passing of RNA molecules through the pore.

**Figure 5.**
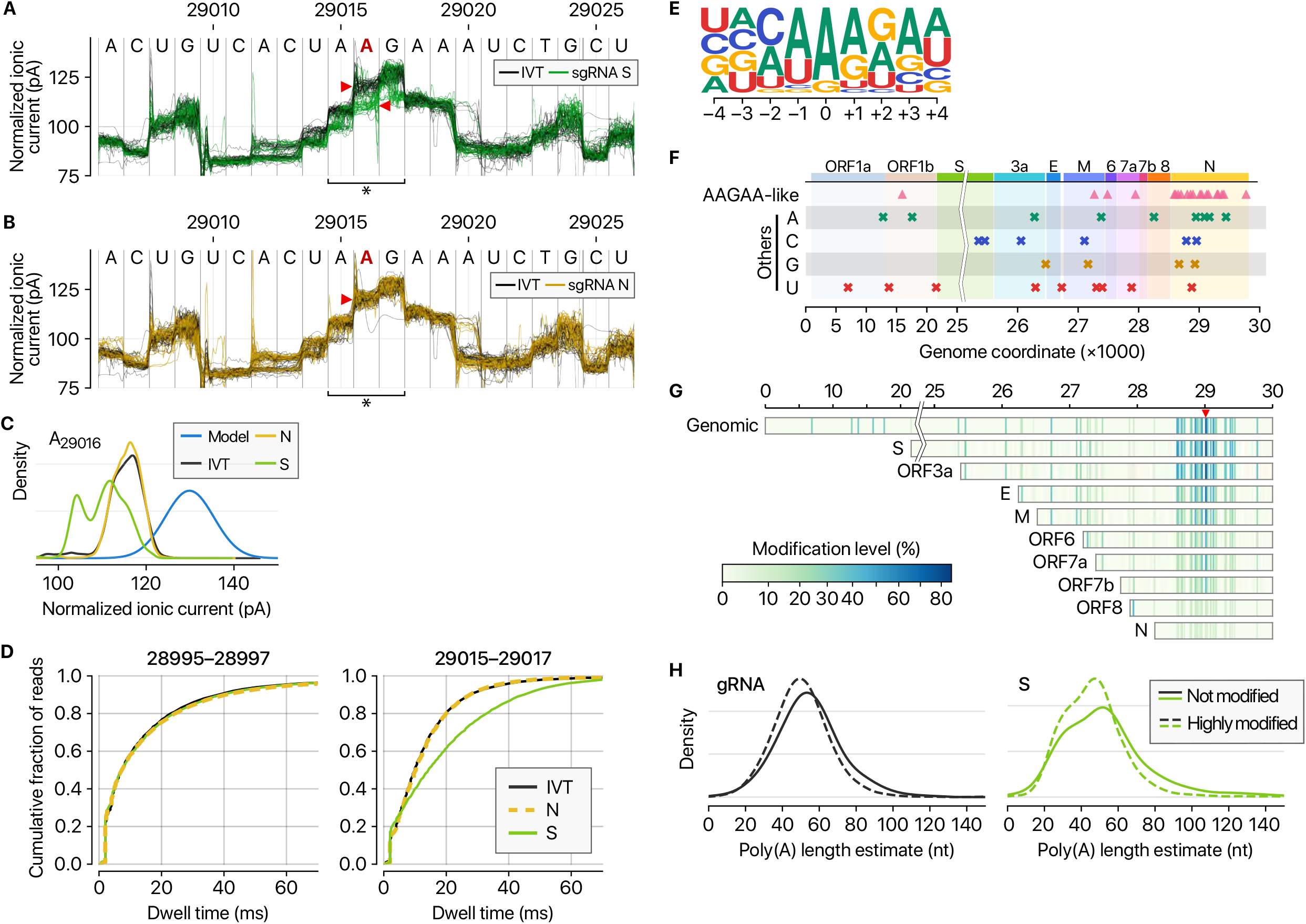
Frequent RNA modification sites. **A,** Distinct ionic current signals (“squiggles”) from viral S transcript (green lines) and in vitro transcribed control (IVT, black lines) indicate RNA modification at the genomic position 29,016. **B,** The ionic current signals from viral N transcript at the genomic position 29,016 (yellow lines) are similar to those from IVT control (black lines), indicating that modification is rare on the N sgRNA. **C,** Ionic current distribution at A29016. Blue line shows the signal distribution in the standard model of tombo 1.5. **D,** Dwell time difference supports RNA modification. The dwell time of the region 29,015-29,017 of the S RNA (right) is significantly longer than those of IVT control and N RNAs. On the contrary, the neighboring region 28,995-28,997 of IVT, N, and S RNA is indistinguishable (left). **E,** Position-specific base frequency of a motif enriched in the frequently modified sites. **F,** Genomic location of modification sites with the AAGAA-like motif (top row) and the others grouped by the detected nucleotide base. **G,** Location and modification levels in different RNA species. **H,** Poly(A) length distribution of gRNA (left) and S RNA (right). Modified viral RNAs carry shorter poly(A) tails.

Among the 41 potential modification sites, the most frequently observed motif is AAGAA (Figures 5E and S2D). The modification sites on the ‘AAGAA-like’ motif (including AAGAA and other A/G-rich sequences) are found throughout the viral genome but particularly enriched in genomic positions 28,500-29,500 (Figure 5F). Long viral transcripts (gRNA, S, 3a, E, and M) are more frequently modified than shorter RNAs (6, 7a, 7b, 8, and N) (Figure 5G), suggesting a modification mechanism that is specific for certain RNA species.

Since DRS allows simultaneous detection of multiple features on individual molecules, we cross-examined the poly(A) tail length and internal modification sites. Interestingly, modified RNA molecules have shorter poly(A) tails than unmodified ones (Figures 5H and S3). These results suggest a link between the internal modification and 3′ end tail. Since poly(A) tail plays an important role in RNA turnover, it is tempting to speculate that the observed internal modification is involved in viral RNA stability control. It is also plausible that RNA modification is a mechanism to evade the host immune response. The type of modification(s) is yet to be identified, although we can exclude METTL3-mediated m6A (for lack of consensus motif RRACH), ADAR-mediated deamination (for lack of A-to-G sequence change in the DNBseq data), and m1A (for lack of the evidence for RT stop). Our finding implicates a hidden layer of CoV regulation. It will be interesting in the future to identify the chemical nature, enzymology, and biological functions of the modification(s).

In this study, we delineate the transcriptomic and epitranscriptomic architecture of SARS-CoV-2. Unambiguous mapping of the expressed sgRNAs and ORFs is a prerequisite for the functional investigation of viral proteins, replication mechanism, and host-viral interactions involved in pathogenicity. An in-depth analysis of the joint reads revealed a highly complex landscape of viral RNA synthesis. Like other RNA viruses, CoVs undergo frequent recombination, which may allow rapid evolution to change their host/tissue specificity and drug sensitivity. The ORFs found in this study may serve as accessory proteins that modulate viral replication and host immune response. The RNA modifications may also contribute to viral survival and immune evasion in infected tissues as the innate immune system is known to be less sensitive to RNAs with nucleoside modification (Kariko et al., 2005). It is yet to be examined if the ORFs and RNA modifications are unique to SARS-CoV-2 or conserved in other coronaviruses. Comparative studies on their distribution and functional significance will help us to gain a deeper understanding of SARS-CoV-2 and coronaviruses in general. Our data provide a rich resource and open new directions to investigate the mechanisms underlying the pathogenicity of SARS-CoV-2.

## Ethics Statement

This study was carried out in accordance with the biosafety guideline by the KCDC. The Institutional Biosafety Committee of Seoul National University approved the protocol used in these studies (SNUIBC-200219-10).

## Acknowledgments

We thank members of our laboratories for discussion and help, particularly Eun-jin Chang and Young-suk Lee. We are grateful to Dr. Jung-Hye Roe and Inhye Park for their advice. We thank Kyung-Chang Kim and Sung Soon Kim at Korea National Institute of Health for their support. The pathogen resource (NCCP43326) for this study was provided by the National Culture Collection for Pathogens, Korea National Institute of Health.

Funding: This work was supported by IBS-R008-D1 of Institute for Basic Science from the Ministry of Science, ICT and Future Planning of Korea (D.K., H.C. and V.N.K.), BK21 Research Fellowship from the Ministry of Education of Korea (D.K.), the New Faculty Startup Fund from Seoul National University (H.C.), and 4845-300(2019-NG044) of Korea Centers for Disease Control and Prevention (J.Y.L., J.S.Y., and J.W.K.).

## Author Contributions

H.C, J.Y.L, and V.N.K. designed the study. D.K., J.S.Y., and J.W.K. performed molecular and cell biological experiments. H.C. carried out computational analyses. H.C., J.Y.L, and V.N.K. wrote the manuscript.

## Accession Numbers

The sequencing data were deposited into the Open Science Framework (OSF) with an accession number doi:10.17605/OSF.IO/8F6N9.

## Supplementary Information

### Supplemental Table Legends

**Supplementary Table 1 | Oligonucleotide sequences used in preparation of IVT control RNAs.**

Refer to Methods for the experimental procedure.

**Supplementary Table 2 | Recombination sites detected from junction-spanning reads.**

The genomic coordinates in the “5′ site” and “3′ site” point to the 3′-most and the 5′-most nucleotides that survive the recombination event, respectively. “Count” is the number of the junction-spanning reads that supports the recombination event. “Skip” is the size of the deletion. “5p_before” is the 3′-most sequence from the 5′-side of the recombination event. “5p_after” is the bases immediately following the “5′ site.” The sequence is the first nucleotides of the region that is excluded by the recombination event. “3p_before” is the last nucleotide sequence of the region that is excluded by the recombination. “3p_after” is the 5′-most nucleotides of the 3′-side of the recombination product.

**Supplementary Table 3 | TRS-L-dependent recombination events resulting in noncanonical leader-body junction (related to Fig. 3E).**

This table extends the information for the recombination events using the canonical leader TRS joined to noncanonical 3′ sites. “Startpos” is the genomic coordinate where the first AUG appears in the resulting recombination product. “Resultingsequence” shows the sequence near the recombination site with the nucleotides from the 5′-side in lowercase, and those from the 3′-side in uppercase. “Annot_cds” is the name of an ORF that shares the start codon position with the recombination product by the “first AUG” rule. “Annot_cds_start” and “annot_cds_end” indicate the interval of the annotated canonical ORF that matches to the start codon in the recombination product. “Startmatch_name” and “startmatch_offset” show the canonical ORF that overlaps with the position of the first appearing AUG in the recombination product, and the distance between the AUG and the start codon position of the ORF. “Stopmatch_name” and “stopmatch_offset” show the canonical ORF that overlaps with the expected stop codon position in case the translation starts at the first appearing AUG, and the relative coordinate of the “expected” stop codon position from the stop codon of the canonical ORF. “Translation” shows the peptide sequence that is expected to be produced by the first ORF in the recombination product.

**Supplementary Table 4 | TRS-L-independent distant recombination events from the 5′ sites located within the “hot spots” (related to Fig. 3F).**

Many distant recombination events do not involve the leader sequence. Instead, the 5′ sites are often found in “hot spots” near ORFs or nsp2 and nsp3. “Count” indicates the number of reads that share the same 5′ and 3′ sites. “Skip” is the size of the deletion in the given combination of 5′ and 3′ sites. “Truncated_prod” is the name of a mature peptide that is truncated by recombination. “Lost_aa” is the number of amino acids lost from the mature peptide by recombination. “Added_aa” is the number of amino acids introduced to the potentially truncated product by recombination. By the recombination event, the sequence indicated in “ctx_orig” turns into the sequence in “ctx_new”.

**Supplementary Table 5 | Detected modification sites in sgRNAs.**

The “pos” column shows all genomic positions reported to be modified at least 20% in any species of sgRNA. “Location” indicates the name of ORF or mature peptide that the position codes. The percentage numbers below the names of sgRNAs are the estimated fraction of modified reads at the position. “Context” shows the sequence context around the modified bases.

**Supplementary Figure 1.**
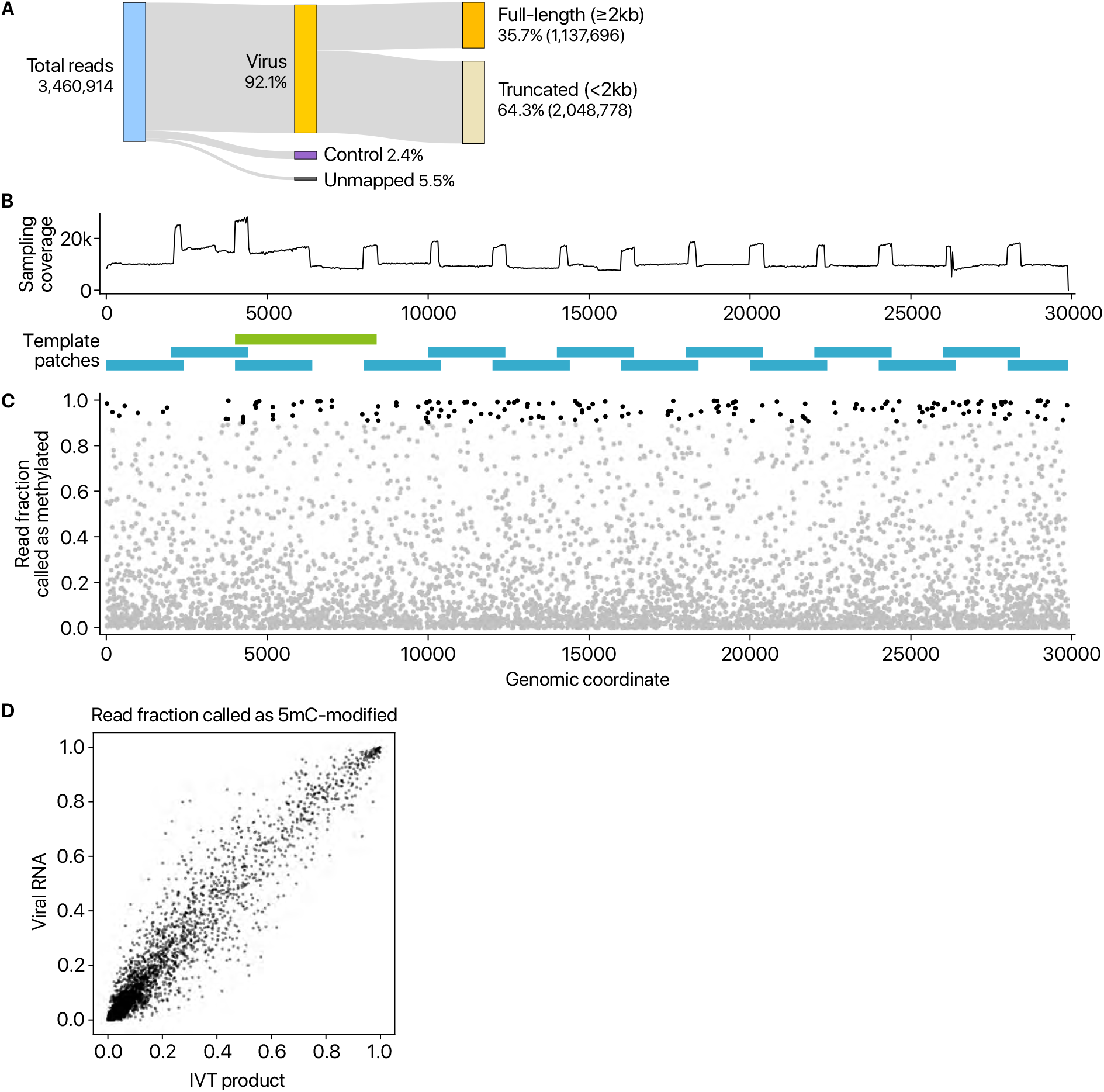
False positive calling of 5mC modification demonstrated by using unmodified negative control RNAs. **A,** Read counts from nanopore direct RNA sequencing of in vitro transcribed (IVT) RNAs that have viral sequences. “Control” indicates quality control RNA for nanopore sequencing. **B,** The 15 partially overlapping patches cover the entire genome (blue rectangles at the bottom). Each RNA is ~2.3 kb in length. One fragment marked with a green rectangle is longer than others (~4.4 kb) to circumvent difficulties in the PCR amplification. The sequenced reads were downsampled so that every region is equally covered. The resulting balanced coverage is shown in the chart at the top. **C,** Detected 5mC modification from in vitro transcribed unmodified RNAs (IVT product) by the “alternative base detection” mode in Tombo. Black dots indicate the sites that satisfy the estimated false discovery rate cut-off calculated using unmodified yeast *ENO2* mRNA (Viehweger et al., 2019). **D,** Comparison between the sites called from unmodified IVT products and those from viral RNAs expressed in Vero cells. **Related to Figure 5**

**Supplementary Figure 2.**
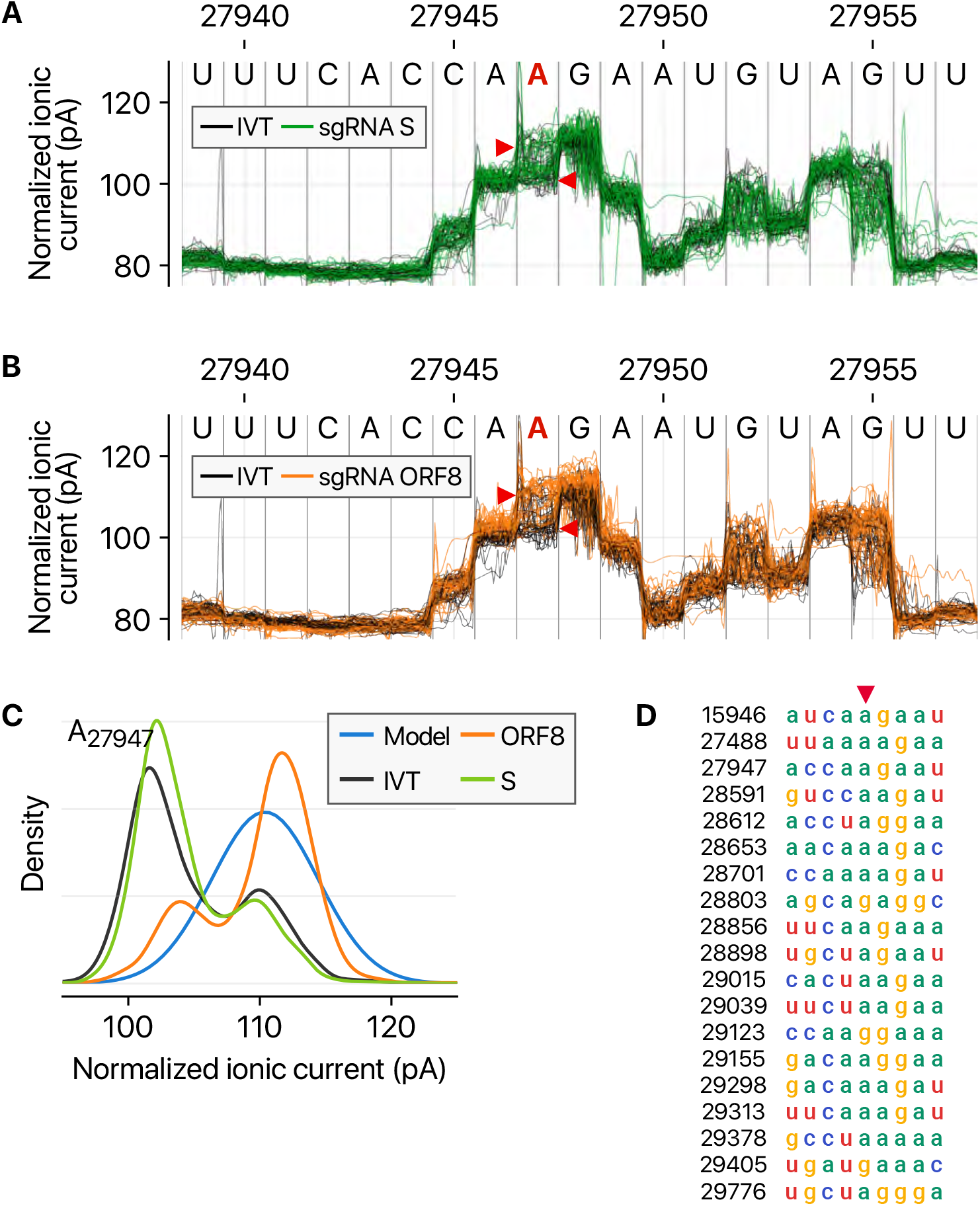
Detected modified sites in viral RNAs. **A,** Ionic current levels near the genomic position 27,947 in viral S RNA (green lines) and IVT control RNA (black lines). **B,** Ionic current levels for the identical region in the viral ORF8 RNA (orange lines) and IVT control RNA (black lines). **C,** Signal distributions at the position 27,947 in the different RNAs. The blue line shows the standard model used for modification detections without controls (“alternative base detection” and “de novo” modes) in Tombo. **D,** Sequence alignment of the detected modification sites with “AAGAA”-type motif. Base positions on the left hand side correspond to the genomic coordinates denoted with red arrowhead. **Related to Figure 5**

**Supplementary Figure 3.**
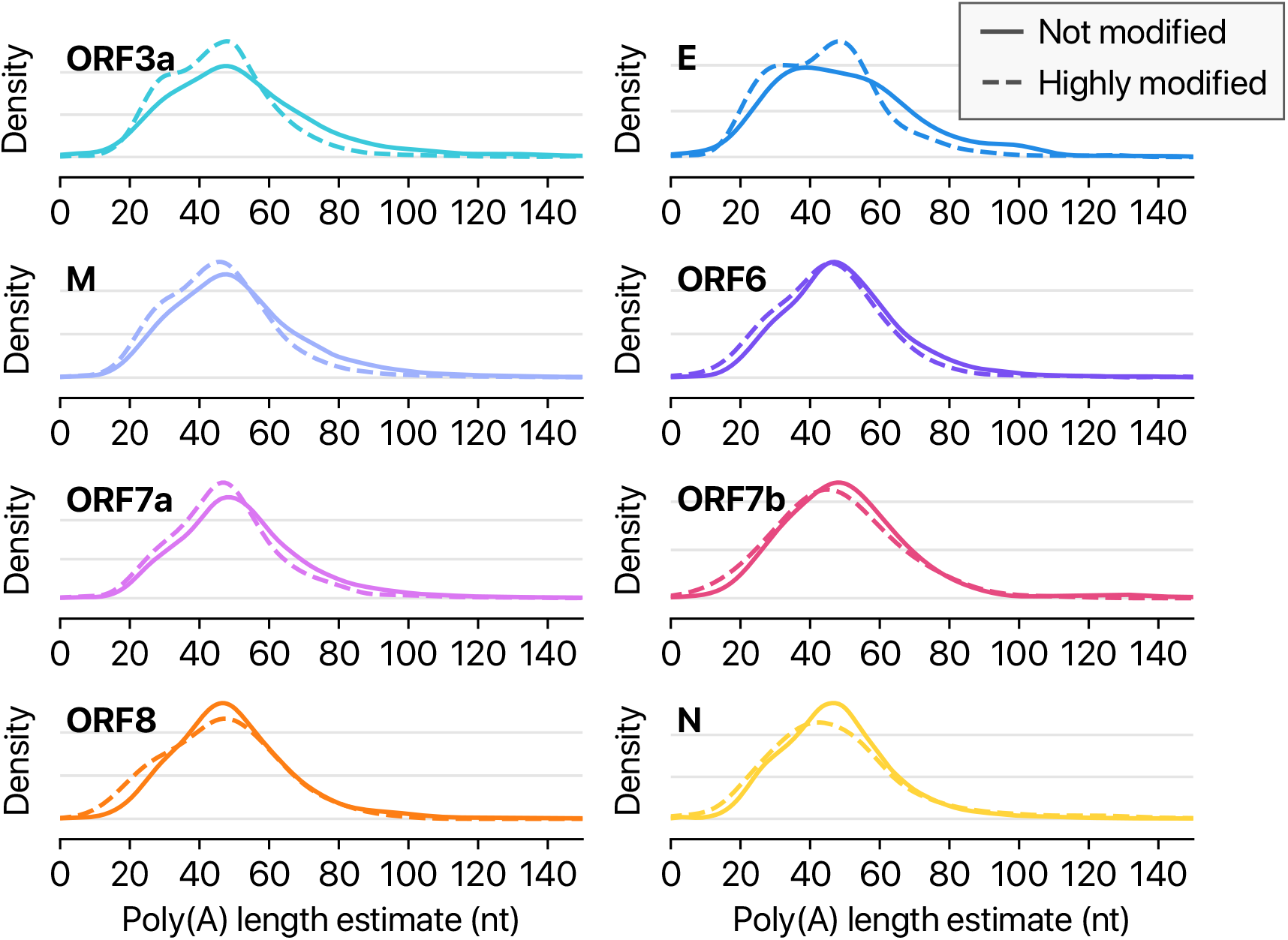
Highly modified viral RNAs carry shorter poly(A) tails. Poly(A) tail length distribution of each viral transcript other than shown in Figure 5. **Related to Figure 5**

## STAR Methods

### SARS-CoV-2 sample preparation

SARS-CoV-2 viral RNA was prepared by extracting total RNA from Vero cells infected with BetaCoV/Korea/KCDC03/2020, at a multiplicity of infection (MOI) of 0.05, and cultured in DMEM supplemented with 2% fetal bovine serum and penicillin-strepamycin. The virus is the fourth passage and not plaque-isolated. Cells were harvested at 24 hours post-infection and washed once with PBS before adding TRIzol. Viral culture was conducted in a biosafety level-3 facility.

### In vitro transcription

Total RNA from SARS-CoV-2-infected Vero cell was extracted by using TRIzol (Invitrogen) followed by DNaseI (Takara) treatment. Reverse transcription (SuperScript IV Reverse Transcriptase [Invitrogen]) was done with virus-specific RT primers. Templates for in vitro transcription were prepared by PCR (Q5® High-Fidelity DNA Polymerase [NEB]) with virus-specific PCR primers followed by in vitro transcription (MEGAscript™ T7 Transcription Kit [Invitrogen]). The oligonucleotides used in this study were listed in Table S1.

### Nanopore direct RNA sequencing

For nanopore sequencing on non-infected and SARS-CoV-2-infected Vero cells, each 4 *μ*g of DNase I (Takara)-treated total RNA in 8 *μ*l was used for library preparation following the manufacturer’s instruction (the Oxford Nanopore DRS protocol, SQK-RNA002) with minor adaptations. 20 U of SUPERase-In RNase inhibitor (Ambion, 20 U/*μ*l) was added to both adapter ligation steps. SuperScript IV Reverse Transcriptase (Invitrogen) was adopted instead of SuperScript III, and the reaction time of reverse transcription was lengthened by 2 hours. The library was loaded on FLO-MIN106D flow cell followed by 42 hours sequencing run on MinION device (Oxford Nanopore Technologies). For nanopore sequencing on SARS-CoV-2 RNA fragments by in vitro transcription, the same method was applied except for a total 2 *μ*g of fragment RNAs and 30 minutes reaction time of reverse transcription. The nanopore direct sequencing data were basecalled using guppy 3.4.5 (Oxford Nanopore Technologies) using the high-accuracy model. The sequence reads were aligned to the reference sequence database composed of the *C. sabaeus* genome (ENSEMBL release 99), a SARS-CoV-2 genome, yeast *ENO2* cDNA (YHR174W), and human ribosomal DNA complete repeat unit (GenBank U13369.1) using minimap2 2.17 (Li, 2018) with options “-k 13 -x splice -N 32 -un”. We used the sequence of the Wuhan-Hu-1 strain (GenBank NC_045512.2) as a backbone for the viral reference genome, then corrected the four single nucleotide variants found in BetaCoV/Korea/KCDC03/2020; T4402C, G5062T, C8782T, and T28143C (GISAID). The sequence alignments were further improved by re-mapping the identified viral reads to the viral genome using minimap2 options “-k 8 -w 1 --splice -g 30000 -G 30000 -A1 -B2 -O2,24 -E1,0 -C0 -z 400,200 --no-end-flt --junc-bonus=100 -F 40000 -N 32 --splice-flank=no --max-chain-skip=40 -un --junc-bed=FILE -p 0.7”. Chimeric reads were filtered out according to the flag from minimap2. The mapped reads from canonical sgRNAs were identified using the start and end positions of large deletions ≥ 10000 nt. For a valid assignment to a species of sgRNA, we required that the start position is between 55 and 85 in the genomic coordinate. The first AUG in the downstream of the end position of a large deletion was used for identification of the “spliced” product. The dwell time of poly(A) tails were measured using poreplex 0.5.0 (https://github.com/hyeshik/poreplex). For the conversion from a dwell time to a nucleotide length, we divided a poly(A) dwell measurement by 1/30 of the mode of the poly(A) dwell time of the ONT sequencing calibration control which has a 30 nt-long poly(A) tail.

### DNBseq RNA sequencing

Total RNA from SARS-CoV-2-infected Vero cell was extracted by using TRIzol (Invitrogen) followed by DNaseI (Takara) treatment. Dynabeads^®^ mRNA Purification Kit (Invitrogen) was applied to 1 *μ*g of total RNA for rRNA depletion. RNA-seq library for 250 bp insert size was constructed following the manufacturer’s instruction (MGIEasy RNA Directional Library Prep Set). The library was loaded on MGISEQ-200RS Sequencing flow cell with MGISEQ-200RS High-throughput Sequencing Kit (PE 100), and the library was run on DNBSEQ-G50RS (paired-end run, 100 × 100 cycles). The sequences from DNBseq were aligned to the reference sequences used in nanopore DRS. We used STAR (Dobin et al., 2013) with many switches to completely turn off the penalties of non-canonical eukaryotic splicing: “--outFilterType BySJout -- outFilterMultimapNmax 20 --alignSJoverhangMin 8 --outSJfilterOverhangMin 12 12 12 12 --outSJfilterCountUniqueMin 1 1 1 1 --outSJfilterCountTotalMin 1 1 1 1 --outSJfilterDistToOtherSJmin 0 0 0 0 --outFilterMismatchNmax 999 --outFilterMismatchNoverReadLmax 0.04 --scoreGapNoncan -4 --scoreGapATAC -4 --chimOutType WithinBAM HardClip --chimScoreJunctionNonGTAG 0 --alignSJstitchMismatchNmax -1 -1 -1 -1 --alignIntronMin 20 --alignIntronMax 1000000 --alignMatesGapMax 1000000”.

### Analysis of junction-spanning reads

The junction-spanning reads (JSR) were prepared by extracting all large deletion (≥ 20 nt) events from the sequence alignments. **[Figure 3A]** We counted the number of 5′ and 3′ sites in 100nt-wide bins. The red dots that indicate the frequently used 3′ sites were identified using the SciPy’s scipy.signals.find_peaks(prominence=2) (Virtanen et al., 2020). **[Figures 3B-3G]** The JSRs were categorized by the position of 5′ and 3′ site positions. A JSR was marked as a leader-to-body junction when the 5′ site of the deletion is between 55 and 85. The sgRNA identity and the frame matching were determined by the first AUG position in the downstream sequence of the 3′ site for the JSRs that the 5′ site is in a region that is annotated as a UTR. For the cases that the 5′ site is in a known ORF or an AUG is introduced by recombination, we checked if the concatenated sequence generates a protein product with the same reading frame as a canonical ORF after the 3′ site.

### Modified base detection and the downstream analyses

The DRS reads of the IVT RNAs were downsampled to balance the coverage between different fragments that were split into equal-sized patches. Sampling frequency of a fragment was controlled by the read counts within a 100-nt bin with the lowest coverage in each fragment. We sampled the reads so that the result contain roughly 10,000 reads from every IVT fragment. The viral RNA reads and the downsampled IVT reads were processed for squiggle analyses by ONT tombo 1.5 (Stoiber et al., 2017) with a minor tune to improve the sensitivity of sequence alignments (-k8 -w1). The modified base detection was done by using the “model_sample_compare” mode with an option “--sample-only-estimates” unless otherwise specified. **[Figure 5E-5F]** The classification of sequence context near the modified sites was first done by the existence of four consecutive purine bases within 5-nt from the position with the highest modification fraction reported by tombo. Then, the rest were further divided into four groups according to the nucleotide base with the highest modification fraction. **[Figures 5H and S3]** “Highly modified” sgRNA reads were detected by referring to eight modification sites which were at least 40% modified in any species of sgRNAs: 28591, 28612, 28653, 28860, 28958, 29016, 29088, 29127. We used the reads that were reported as modified at three or more sites with a statistic < 0.01 as “highly modified” reads. “Not modified” reads were reported with the statistic ≥ 0.01 in all eight sites.

